# CGMD: An integrated database of Cancer Genes and Markers

**DOI:** 10.1101/011262

**Authors:** Pradeep K Jangampalli Adi, Kranthi K Konidala, Nanda K Yellapu, L Balasubramanyam, Bhaskar Matcha

## Abstract

Cancer is a dysregulation of apoptosis process a programmed cell death through which the cell number is tightly regulated. Many factors affect apoptosis including tumor genes (TGs). Moreover, several researchers identified TGs and demonstrated their functions in various types of tumors or normal samples. Surprisingly, it has also been indicated that the expression of tumor markers in the development of cancer type is also one of the important target sites for clinical applications. Therefore, integrating tumor genes and tumor markers with experimental evidences might definitely provide valuable information for further investigation of TGs and their crosstalk in cancer. To achieve this objective, we developed a database known as Cancer Gene Marker Database (CGMD) which integrates cancer genes and markers based on experimental background. The major goal of CGMD is to provide: 1) systemic treatment approaches and advancements of different cancer treatments in present scenario; 2) Pooling of different genes and markers with their molecular characterization and involvements in different pathways; 3) Free availability of CGMD at www.cgmd.in. The database consists of 309 genes plus 206 markers and also includes sequences of a list of 40 different human cancers and moreover, all the characterized markers have detailed descriptions in appropriate manner. CGMD extraction is collective and informative by using web resources like National Cancer Institute (NCI) and National Center for Biotechnology Information (NCBI), UniProtKB, KEGG and EMBOSS. CGMD provides updated literature regarding different cancer annotations and their detail molecular descriptions such as CpG islands, promoters, exons of cancer genes and their active sites, physico-chemical properties and also systematically represented the domains of characterized proteins.

---

Database URL: http://cgmd.in/

## Introduction

Cancer is treated as a hallmark of abnormal cell proliferation which eventually leads to an imbalance between viable and death cells. It is well accepted that genetic changes are one of the important causes for cancer (Fulda et al.2010). Though, the exact root cause for these genetic changes is not well defined, it has been suggested that mutations or alterations in the chromosome complement at least in part leads to changes in genome architecture thereby leads to cancer (International Human Genome Sequencing Consortium 2004). Recent evidences also indicated that mutations in more than 1% of genes contribute to human cancer (Futreal et al.2004) Moreover, these mutations are responsible for switching of proto-oncogenes to oncogenes and/or loss of function in case of tumor suppressor genes. Proto-oncogenes are the normal genes which code for proteins that regulate the process of cell proliferation differentiation and apoptosis (Todd and Wong 1999) and thus, even a small change due to point mutations and/or chromosomal aberrations such as translocations, duplications, and deletions convert the normal genes into oncogenes. It has been also claimed that mutations occurring in regulatory regions especially in the promoter sequences and/or epigenetic mechanisms (Renan 1993) are the primary root causes for mutation type cancers. Mutations at the level of promoter regions switches the gene regulatory mechanisms and epigenetic changes such as hypo-or hyper-methylation processes may lead to chromosomal instability, altered expression and transcriptional silencing of tumor suppressor genes that may lead to the development of tumors (Esteller 2002). One of the recent findings indicated that non-protein-coding RNAs can act as tumor suppressor genes thereby control cell proliferation and apoptosis at the post-transcriptional level during neoplasm development (Andreeff et al.2000). It appears that mutations target genes involved in cell-proliferation regulatory machinery system thereby leads to cancer. Therefore, understanding the basic roots of cancer at cellular and molecular level i.e., genes, their translated products (proteins) and their role in biochemical mechanisms might provide valuable insights.

Many research groups by using high-throughput strategies showed that expression of tumor suppressor genes changes with respect to cancer stage and type. It is evident that in addition to tumor genes, data related to tumor markers provide signals to get full picture of cancer stage and type. Thus, tumor markers will provide invaluable clues for identifying genes highly expressed in tumors but not in normal adult tissue. However, the freely available resource databases dealing with both tumor genes and markers are yet to be developed. Therefore it is prime concern, integrating the available experimental data related to tumor genes at the level of genome, transcriptome and proteome levels is considered important for fully understanding the alterations in genes. To accomplish this task, a systematic integration of experimental evidence is required to develop which carefully catalog known cancer genes and markers from accumulated diverse literature and evaluate their consistency. Earlier, there were two databases available such as TSGDB (Yang and Fu 2003) and TSgene (Zhao et al. 2003) which contributed to integrate cancer related works based experimental evidences. TSGDB is a database developed based on only experimental evidences and on the other hand, TSgene is a comprehensive database which taken all resources from UniportKB (The UniProt Consortium 2012), the Tumor Associated Gene (TAG) database and PubMed abstracts. Therefore, an attempt has been made to develop a resource database known as cancer gene marker database (CGMD) which includes information related to both tumor suppressor genes and tumor markers by integrating experimental evidences (http://www.ncbi.nlm.nih.gov/pubmed/) and cancer related web-based tools. CGMD consists of the literature supported data from well-established databases like KEGG genes (Samuelsson et al.2010) NCBI catalogs (Vogelstein and Kinzler 2004) and UniProtKB (The UniProt Consortium 2012). It covers a broad range of literal information and advancement of treatments in cancer and updated information of characterized cancer genes and markers at genomic and proteomic levels with large scale experimental evidences. Furthermore, we manually collected and curated in a systematic way to integrate experimental data through high throughput screening in CGMD (http://www.cgmd.in) In this current version of CGMD provides 309 cancer genes and 206 markers totally 515 genes, corresponding to 40 different cancers genes and markers information. Moreover, CGMD is encircling different databases to encompass information related to cancer genes. For example Kyoto Encyclopedia of Genes and Genomes (KEGG) entry, position of the gene, CpG islands, promoter regions, exons of different cancers and markers at molecular and proteomic levels should be used. For accuracy quick calling response for the user we developed a MySQL database which will be helpful to the user in interface mode to access the data. The fundamental objective of CGMD database is to collect and integrate cancer genes and markers from the different database resources and to provide elaborative and detailed information of annotated cancer genes and markers. This database simply provides corresponding cross-linking and exact query information with user interface. This freely developed database is user friendly and at the same time acts as a platform for better understanding the molecular futures of cancer genes and markers by giving the reliable information and helpful for cancer research communities and also to medical oriented diagnostic developments.

## Mining for Data

The data related to cancer genes and markers were collected from public databases such as NCBI, UniProt KB, KEGG disease and EMBOSS and organized the collected data systematically. Initially, we listed different types of cancers from NCI-PDQ, web resource (Kanehisa et al.2004) then queried each cancer marker sequence through Kyoto Encyclopedia of Genes and Genomes (KEGG) and eventually characterized the sequences. All the sequences were checked with PubMed literature evidence. Later, manual analysis process was performed to analyze data pool at molecular level through online bioinformatics tools. Moreover, CGMD to provide reliable and detailed extensive information related to cancer markers, Entrez and PubMed biomedical literature search system and online sequence analysis tools provides the analyzed data and sequentially organized in advanced information of cancer genes and markers.

**Table 1.**
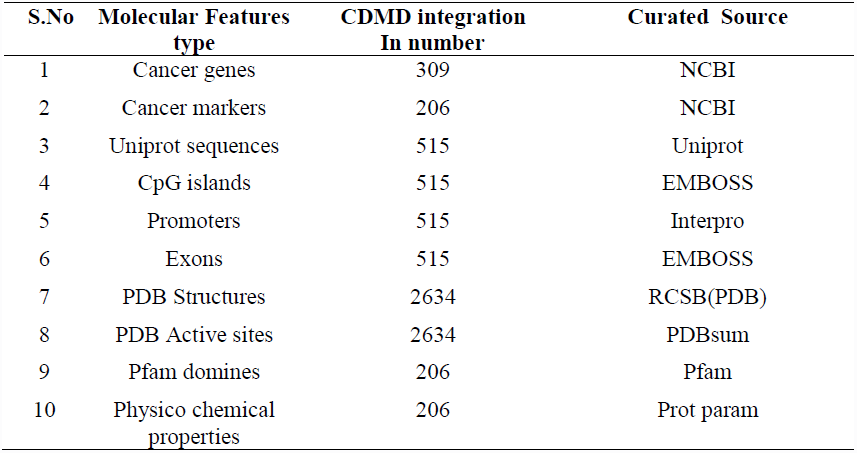
The CGMD complete data of molecular features of cancer genes and markers and online analysis data and curation sources tabulated.

Categorization of data in CGMD broadly divided in to two major concerns like one is for genes and another one for markers. Generally there is no difference between genes and markers but CGMD differentiate the mutated whether it may be germ line or somatic based and each characterized mutation can be considered as each mutation a marker and the corresponding sequence are the gene. For example BRAC1 gene contain 3 mutations here the BRAC1 is the cancer gene and the each 3 characterized mutations will be considered as 3 independent marker sequences. Firstly we manage to align the cancer genes by searching the respective keywords in the NCBI from the pool of sequences, we isolated the respective sequences IDs through KEGG disease and short listed to gives an accurate data of cancer genes. Each Entrez IDs are listed separately and cross checked with PubMed abstracts, care has been taken to not overlap with the cancer gene sequences and with markers sequence. An individual unique both cancers genes and marker genes are prepared and sets next step for the analysis. Each cancer sequence IDs taken as the key element to retrieve the information of both cancer genes and markers in various genomic and proteomic tools. These analyzed output data again cross checked with raw sequence of cancer genes and markers to minimize the error prone. After checking of all these needful things we manually draft a separate genomic and proteomic data for both cancer genes and markers. Finally we pool the information nearly 2265 of Promoters and exons and of six frame translation products of each cancer gene. In proteomic approach we managed to list the all possible 2634 PDB structures from KEGG disease and domain descriptions and corresponding SNPs etc. For better understanding of functional annotations of cancer genes and markers in CGMD database we provide the basic information as well as the extensive functional futures of genes and markers are tabulated (Table 1). Basic information of cancer genes through UniProt entries and cross linking databases like PDBsum provides the core futures of cancer proteins (National Library of Medicine 2013). We made extended interlinking with KEGG Pathway database to know the cancer markers functional pathways (NCI 2013). And finally to know the superfamily characteristics of cancer markers should be known by Pfam database (Finn et al.2008).

## Significant Database futures

Over all data among the cancer markers and genes about 453 CpG islands were detected known the conserved regions of the sequence that can be protected incidental errors in genome (Illingworth et al.2010). Then the promoters, the regulatory sites distinct regions these are the functional switches of genes may be turned on or off by tuning the structural features characteristic for each chromosomes (Ong and Corces 2011). Exon regions are gene predicting evidences to known from where exactly the carcinogenic sequence turns to pathological protein products, which helps for prediction of cancer protein sequences in all possible frames (Olivier et al.2010). BLAST feature provides quick research with corresponding cancer genes and marker with data base sequences. Data base provides the PDB structures and PDB active sites by crosslinking with PDB and PDBsum database (Laskowski et al.1997) CGMD analyses the proteins physico chemical properties, like molecular weight, pI, amino acid composition, Instability index where provides the protein stability and aliphatic and grand average of hydropathicity (GRAVY) of cancer genes and markers through Prot param tool (Hosseinzadeh et al. 2012). The detailed contents of the CGMD illustrated in (Figure 1).

**Figure 1.**
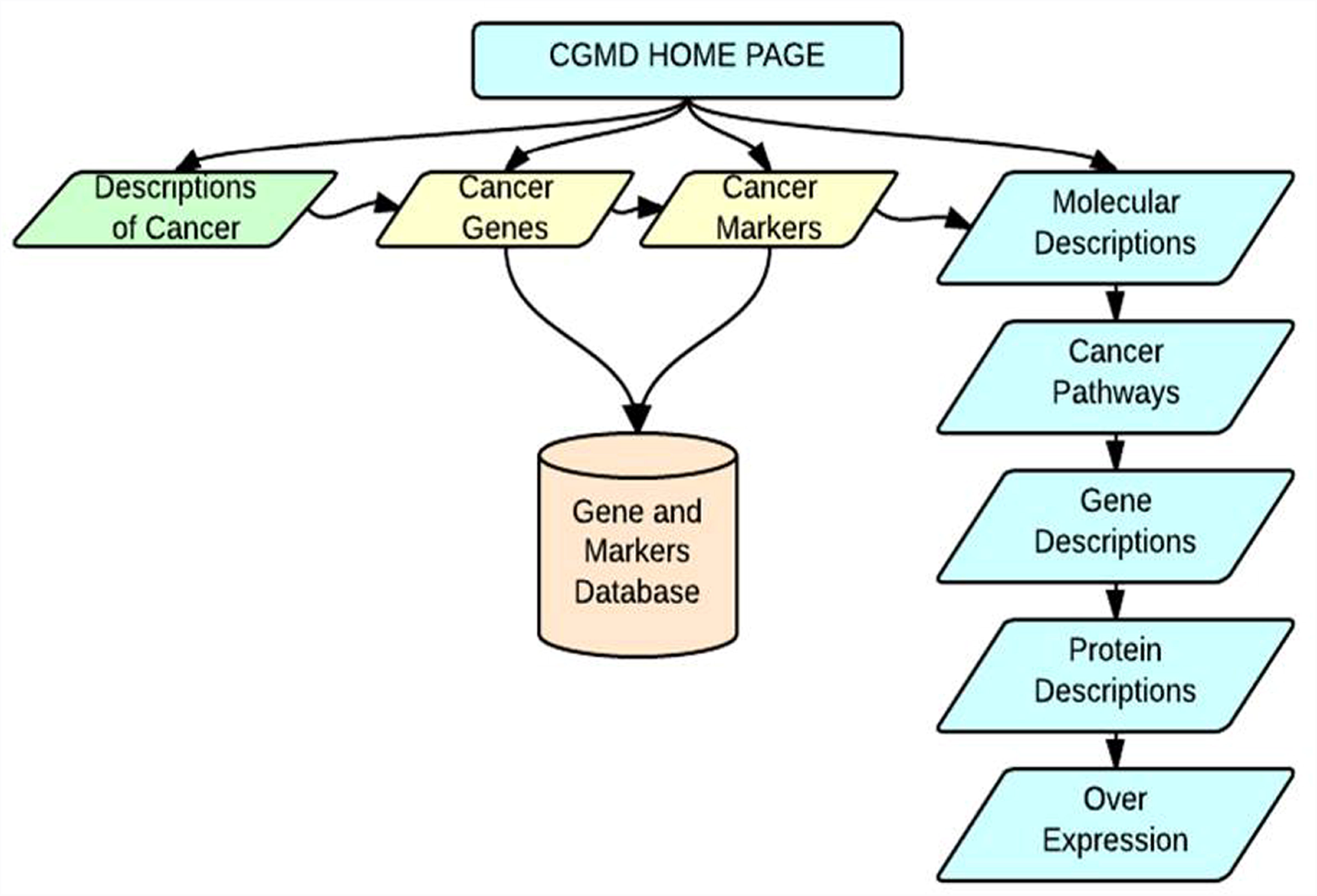
Workflow for processing and analyzing of genomic and proteomic sequences of different human cancers and in-depth transcriptomic sequences specific to different human cancer types.

## Design and Implementation

We designed a MySQL database to store the complete data and front end tool PHP will interface user to get data and browsing. Database infrastructure enabling efficient storage and retrieval, and database design to accommodate different resources to communicate with each other, and to allow users to access the information through PhpMyAdmin and JavaScript. An overview of the database architecture is shown in (Figure 2).

**Figure 2.**
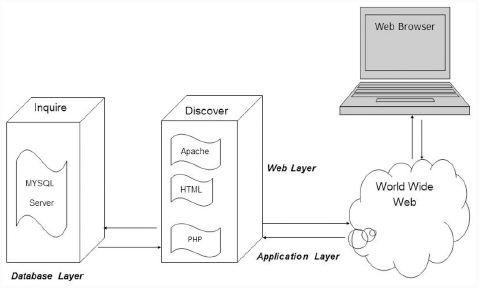
Overview of CGMD architecture.

## Key entries in CGMD

The comprehensive CGMD database homepage shown in (Figure 3A). Home page having the different tabs ‘Description,’ ’Genes,’ ‘Markers,’ Molecular descriptions,’ and each tab labeled separately to represent the stored information related to the concern data. By clicking the tab ‘Description,’ display the list of cancers alphabetically from A to Z by selecting particular alphabet and it displays the list of cancers recorded under the particular letter, (Figure 3B&C), the list was prepared by the updated information National Cancer Institute (NCI) (Kanehisa et al. 2013). In this tab represents the general information of cancers like disease information, symptoms, pathogenicity, advances in treatments and finally updated literature evidence. The second tab of the database which is literally supported data appears in the ‘Genes’ tab nearly 309 number of genes included for 40 types of cancers and the respective sequences are stored (Figure 3D&E). In the next tab that is ‘Markers’ we have collected almost 206 markers for A to Z cancers and their protein sequences are sequentially listed. Finally we encounter (reached) for the important prospects of CGMD that is ‘Molecular descriptions’ under this tab we included the ‘pathways,’ ‘gene description,’ ‘protein description,’ (Figure 3F&G) by clicking the ‘pathways’ it representing by displaying the list of cancer markers and its pathways and the number of tumor suppressor genes involvement and their detailed descriptions with respective types of cancers (Figure 3H). By clicking the gene descriptions it displays the new navigation window which is included the total listed cancers and their markers which help to known the detail information of cancer gene (Figure 3I). By clicking on the Gene ID number it displays the preloaded annotated analyzed data and crosslinking detailed information of genes (Figure 3I). If user wants to know the markers information the only thing to click Marker ID number. For ‘protein description’ the new navigation window displayed with preloaded data of protein information for the respective markers will be appeared by clicking the Marker ID number (Figure 3J&K). Finally we made a simple bar chart for over expressed marker and genes (Figure 3L).

**Figure 3.**
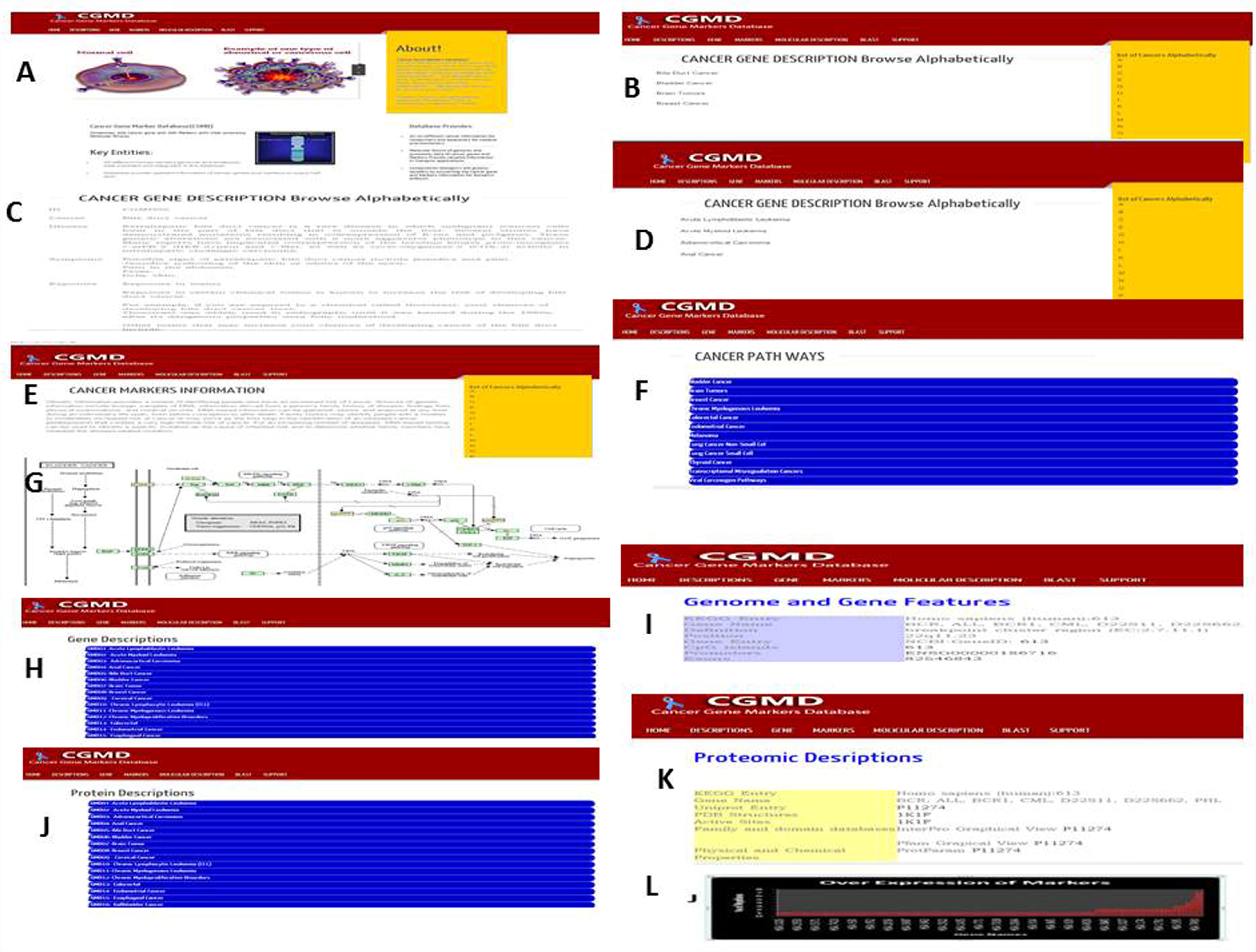
Web interface of the CGMD database. (**A**) Basic home page diplayingthe information in the CGMD database. (**B**) Different Cancer information display list from A to Z. (**C**) A typical highlighted literature with supporting keywords browses by alphabetically. (**D**) Gene description profile. (**E**) Query interface. (**F**) Browser for various cancer pathway types. (**G**) KEGG pathway mapped with CGMD (color-marked). (**H**) Browsing gene description list. (**I**) Provide genome and gene features. (**J**) Browsing description list types (protein-coding). (**K**) Provide proteomic description features. (**L**)Over expressed markers list in CGMD database.

## BLAST

To help users perform sequence alignment, BLASTP was embedded in the database as using this BLAST tab (www.Blastp program (v2.2.26) to provide a graphic user interface through the Web forms and Users can perform similarity searches against each type of various cancer sequences using BLASTP.

## Inference about CGMD

CGMD resource tool is user friendly. The resource tool includes tumor genes and tumor markers with a particular identity (ID) codes which are intended to clearly distinguish genes and markers. Moreover, the CGMD-IDs have been interlinked with Entrez ID codes which pave a way to genome location from original literature source. Interlinking the web resources for cancers genes and markers and extracting the online tools output data form the database is manageable to user.

## Availability and Future Directions

CGMD is a free available resource database which provides information related to cancer genes and markers by integrating several factors including genomic, proteomic, methylation and interactions of predefine active sites. The rationale for the development of CGMD is to provide a valuable resource database for better understanding tumor genes at molecular level which will be useful for therapeutic applications. Finally, we will continuously collect and curate tumor genes and markers by integrating experimental evidences and update CGMD information for understanding full picture of tumor related pathways.

## Abbreviations

TSGDB: tumor suppressor gene database; TAG: Tumor Associated Gene; KEGG: Kyoto Encyclopedia of Genes and Genomes; EMBOSS: European Molecular Biology Open Software Suite; NCI-PDQ: National Cancer Institute Physician Data Query; PDB: Protein data bank; CGMD: cancer gene markers database.

## Authors’ contributions

PKJA developed the original concept, implemented the demonstration version, designed the website template, and wrote the majority of the manuscript. KKK helped in testing the completed tool and gave suggestions for additional query options. NKY and BL designed the database and built the web application's front end. BL developed the demonstration version and wrote the query and data format scripts. BM directed the molecular profiling project, helped write the manuscript, and made input at every step of the database development. All authors read the final manuscript.

## Acknowledgements

We highly thankful to DBT BIF lab for providing infrastructure lab to make this work and I deeply convey my best regards with our lab mates who encourage to carrying out this work.

### Funding

None

## Reference

Andreeff M, Goodrich DW, Pardee AB. 2000. Cell proliferation, differentiation, and apoptosis, in Holland-Frei Cancer Medicine, Section 1 Cancer Biology, BC Decker, 5th edition, chapter 2, pp. 17–32.

Esteller M. 2002. CpG island hypermethylation and tumor suppressor genes: a booming present, a brighter future. Oncogene 21:5427–5440.

Finn RD, Tate J, Mistry J. 2008. The Pfam protein families database. Nucleic Acids Res 36: D281–D288.

Fulda S, Gorman AM, Hori O, Samali A. 2010. Cellular Stress Responses: Cell Survival and Cell Death. Int J Cell Biol 214074. doi: 10.1155/2010/214074.

Futreal PA, Coin L, Marshall M, Down T, Hubbard T, Wooster R, Jones ML, Thornton JM. 2004. A census of human cancer genes. Nat Rev Cancer 4:177–183.

Hosseinzadeh F, Ebrahimi M, Goliaei B, Shamabadi N. 2012. Classification of Lung Cancer Tumors Based on Structural and Physicochemical Properties of Proteins by Bioinformatics Models. PLoS ONE 7:e40017.

Illingworth RS, Gruenewald-Schneider U, Webb S. Kerr AR, James KD, Turner DJ, Smith C, Harrison DJ, Andrews R, Bird AP. 2010. Orphan CpG islands identify numerous conserved promoters in the mammalian genome. PLoS genetics 9: e1001134.

International Human Genome Sequencing Consortium. 2004. Finishing the euchromatic sequence of the human genome. Nature 43:931–945.

Kanehisa M, Goto S, Kawashima S, Okuno Y, Hattori M. 2004. The KEGG resource for deciphering the genome. Nucleic Acids Res 32: D277–D280.

Laskowski RA, Hutchinson EG, Michie, AD. Wallace AC, Jones ML, Thornton JM. 1997. PDBsum: a Web-based database of summaries and analyses of all PDB structures. Trends Biochem. Sci 22: 488–490.

National Library of Medicine. Pub Med. Available at: Febrauray 9, 2013. [http://www.ncbi.nlm.nih.gov/sites/entrez].

NCI comprehensive cancer database available at [http://cancer.gov/cancertopics/pdq] Accessed on 02, 03-2013.

Olivier M, Hollstein M, Hainaut P. 2010. TP53 mutations in human cancers: origins, consequences, and clinical use. Cold Spring Harb Perspect Biol 2: a001008.

Ong CT, Corces, VG. 2011. Enhancer function: new insights into the regulation of tissue-specific gene expression. Nat Rev Genet 12: 283–93.

Renan MJ. 1993. How many mutations are required for tumorigenesis? Implications from human cancer data. MolCarcinog 7:139–146.

Samuelsson JK, Alonso S, Yamamoto F, Perucho M. 2010. DNA fingerprinting techniques for the analysis of genetic and epigenetic alterations in colorectal cancer. Mutat Res 693:61–76. doi: 10.1016/j.mrfmmm.2010.08.010.

The UniProt Consortium: 2012. Reorganizing the protein space at the Universal Protein Resource (UniProt). Nucleic Acids Res 40: D71–D75.

Todd R, Wong DT: Oncogenes. 1999. Anticancer Res 19:4729–4746.

Vogelstein B, Kinzler KW. 2004. Cancer genes and the pathways they control. Nature Medicine 10: 789–799.

Yang Y, Fu LM. 2003. TSGDB: a database system for tumor suppressor genes. Bioinformatics 19:2311–2312.

Zhao M, Sun J, Zhao Z. 2013. TSGene: a web resource for tumor suppressor genes. Nucleic Acids Res 41: D970–D976.

